# m^6^A methylated long noncoding RNA regulates proinflammatory response emerging as novel target for IBD

**DOI:** 10.1101/2023.01.17.524371

**Authors:** Ane Olazagoitia-Garmendia, Henar Rojas-Márquez, Maialen Sebastian-delaCruz, Anne Ochoa, Luis Manuel Mendoza-Gomez, Alain Huerta Madrigal, Izortze Santin, Ainara Castellanos-Rubio

**Author notes:** Corresponding author Twitter: @FunImmune, @AinaCastellanos.

## Abstract

Cytokine mediated sustained inflammation increases the risk to develop different complex chronic inflammatory diseases, such as inflammatory bowel disease (IBD). Recent studies highlighted the involvement of inflammation associated gene variants in m^6^A methylation. Moreover, long noncoding RNAs (lncRNAs) participate in the pathogenesis of inflammatory disorders and their function can be influenced by differential methylation. Here we describe the functional implication of *LOC339803* lncRNA in the development of IBD. We found that allele-specific m^6^A methylation affects YTHDC1 mediated protein binding affinity. *LOC339803*-YTHDC1 interaction dictates chromatin localization of *LOC339803* ultimately inducing *IL1B* and contributing to the development of intestinal inflammation. Our findings were confirmed using human intestinal biopsy samples from IBD and controls.

Overall, our results support *LOC339803* lncRNA as an important mediator of intestinal inflammation, presenting this lncRNA as a potential novel therapeutic target for the treatment of IBD.

## INTRODUCTION

Chronic inflammatory diseases are a wide range of autoimmune and inflammatory diseases characterized by persistent inflammation. Specifically, intestinal inflammatory disorders are a group of diseases in which inflammation is present along the gastrointestinal (GI) tract. There are several inflammatory diseases connected to the digestive system, being inflammatory bowel disease (IBD), comprised of ulcerative colitis (UC) and Crohn’s disease (CD), one of the most common (*1*). In IBD, aberrant cytokine responses to environmental triggers (viral infections, microbiota dysbiosis, dietary agents, etc.) produced not only by the immune cells but also by non-immune cells, such as epithelial cells, lead to chronic inflammation and destruction of healthy tissues. The sustained inflammation results in intense abdominal pain and diarrhea and it can also lead to serious intestinal and extraintestinal complications, as GI cancer or psychological symptoms (*2, 3*). Indeed, people with IBD are at an increased risk of developing GI cancer, particularly colon cancer (*4–6*). Even if there are several therapeutic strategies used to treat IBD, including different types of medication, surgery, and lifestyle changes, there is no actual cure for this disease, highlighting the need of more effective and specific treatments (*7*).

Genetic and environmental factors have been identified to play a major role in the development of these disorders. While Genome Wide Association Studies (GWAS) and Immunochip studies have helped to identify *loci* that confer risk to these pathologies, the understanding of their underlying mechanism remains limited, mostly due to their localization in noncoding genomic regions and their individual small effect size (*8, 9*). Within the last decade, advances in RNA sequencing techniques have revealed novel noncoding RNAs, from which a high number corresponds to the family of long noncoding RNAs (lncRNAs) (*10, 11*). LncRNAs are RNA molecules longer than 200 nucleotides with no or low protein coding potential. They have been involved in key cellular processes through a wide diversity of mechanisms, as they are able to bind DNA, RNA or proteins. Indeed, lncRNAs can respond to external stimuli in a quick and cell type-specific manner, working both transcriptionally and post-transcriptionally (*11–13*). Interestingly, inflammation-associated single nucleotide polymorphisms (SNPs) are enriched in lncRNAs (*12*) and some lncRNAs have been associated with IBD (*14–16*). Hence deciphering the role of inflammation associated SNPs in lncRNA function may help to understand the pathogenesis of this complex disorder.

Some other works have also highlighted that inflammation-associated SNPs can affect RNA modifications as N^6^-methyladenosine (m^6^A) (*17, 18*). m^6^A is the most abundant internal chemical modification of mRNAs and noncoding RNAs and it is involved in multiple aspects of RNA metabolism, playing crucial roles in many cellular processes. In the last years, research on m^6^A-mediated regulatory pathways has increased exponentially (*19–22*) and recent work has suggested that m^6^A may be involved in the development of IBD (*23–25*) . Even if m^6^A-quantitative trait loci (QTL) have been described to be expression and splicing QTL independent (*17*), little is known about the genetic effects of m^6^A modification and their role in diseases. Considering that more than 85 % of inflammation-associated SNPs are located in noncoding regions and have difficult to assess functions, new approaches are needed to clear up the biological functions of noncoding SNPs. Addressing functional characterization studies of these variants by integrating m^6^A methylation data may help to identify new regulatory effects of disease-specific associated SNPs, opening the door to the development of much needed novel therapeutic approaches.

In this work, we studied the lncRNA *LOC339803*, a lncRNA with a previously unknown function located on the Immunochip region 2p15. This region is associated with several inflammatory disorders including IBD. Interestingly, the SNP rs11498 is located next to a m^6^A motif making *LOC339803* an interesting candidate to study the effect of differential m^6^A methylation on lncRNA function in the context of intestinal inflammation. Here, we show that *LOC339803* presents allele-specific m^6^A methylation levels in intestinal cells, affecting YTHDC1 mediated protein binding affinity. YTHDC1 interaction with *LOC339803* activates the nuclear factor kappa B (NFκB) key inflammatory pathway, which will ultimately cause higher basal *IL1B* inflammatory cytokine levels in the individuals with the risk allele. Moreover, we also demonstrated that targeting either *LOC339803* or its m^6^A motif has the ability to ameliorate inflammation.

Overall, our results reveal that m^6^A dependent *LOC339803* lncRNA induces *IL1B* cytokine and present a functional characterization of an inflammation associated SNP, explaining its implication in the development of IBD and opening the door to novel therapeutic interventions.

## RESULTS

### rs11498 SNP genotype affects m^6^A methylation levels and stability of *LOC339803* lncRNA

The SNP rs11498 (GRCh38:chr2:61,143,684), within the IBD-associated Immunochip region 2p15, is located within an exon of the uncharacterized lncRNA *LOC339803* (also known as *AC016747.3*), close to an m^6^A methylation motif (Fig. 1A). This region is associated to various inflammatory disorders including multiple sclerosis, psoriasis, rheumatoid arthritis and both forms of IBD, Crohn’s’ disease and ulcerative colitis, (CRO and UC respectively) (Fig. S1A). Considering that lncRNAs generally have cell-type specific expression, we first quantified *LOC339803* expression in a commercially available RNA pool of different human tissues. As it has been previously observed in many other lncRNAs, *LOC339803* was differentially expressed among the analyzed tissues, showing lowest expression levels in GI tissues as colon or small intestine and highest in the brain (Fig. S1B). RNAseq data retrieved from GTEx database (*26*) also showed similar tissue specific expression patterns (Fig. S1B). Thus, the differential expression levels observed among tissues suggest that this lncRNA could have cell type specific functions contributing to the development of different inflammatory diseases. Interestingly, and supporting this idea, while the G allele confers risk for intestinal inflammation, the opposite allele, A, is the risk allele for other organ specific immune disorders such as multiple sclerosis or psoriasis. An examination of the MetDB m^6^A database (*27*) revealed m^6^A peaks around the SNP (Fig S1.C) suggesting that the region surrounding the SNP could be in fact methylated. Moreover, using online m^6^A predictor SRAMP (*28*), we observed that the m^6^A motif located next to the SNP presents a high probability of methylation (Fig. S1D) and the predicted secondary structure of the site changes depending on the rs11498 genotype, with the G allele form showing a more accessible site (Fig. 1B). In order to confirm these predictions, we used the HCT-15 intestinal epithelial cell line as an *in vitro* model of IBD, as it is heterozygous for the SNP rs11498. METTL3 RNA immunoprecipitation (RIP) confirmed the binding of the m^6^A writer to *LOC339803* (Fig. 1C), indicating that the m^6^A motif next to the associated SNP could be indeed methylated. Moreover, allele-specific meRIP in the intestinal cell line further confirmed the methylation of the site, with the G allele being preferentially methylated (Fig. 1D).

**Fig. 1.**
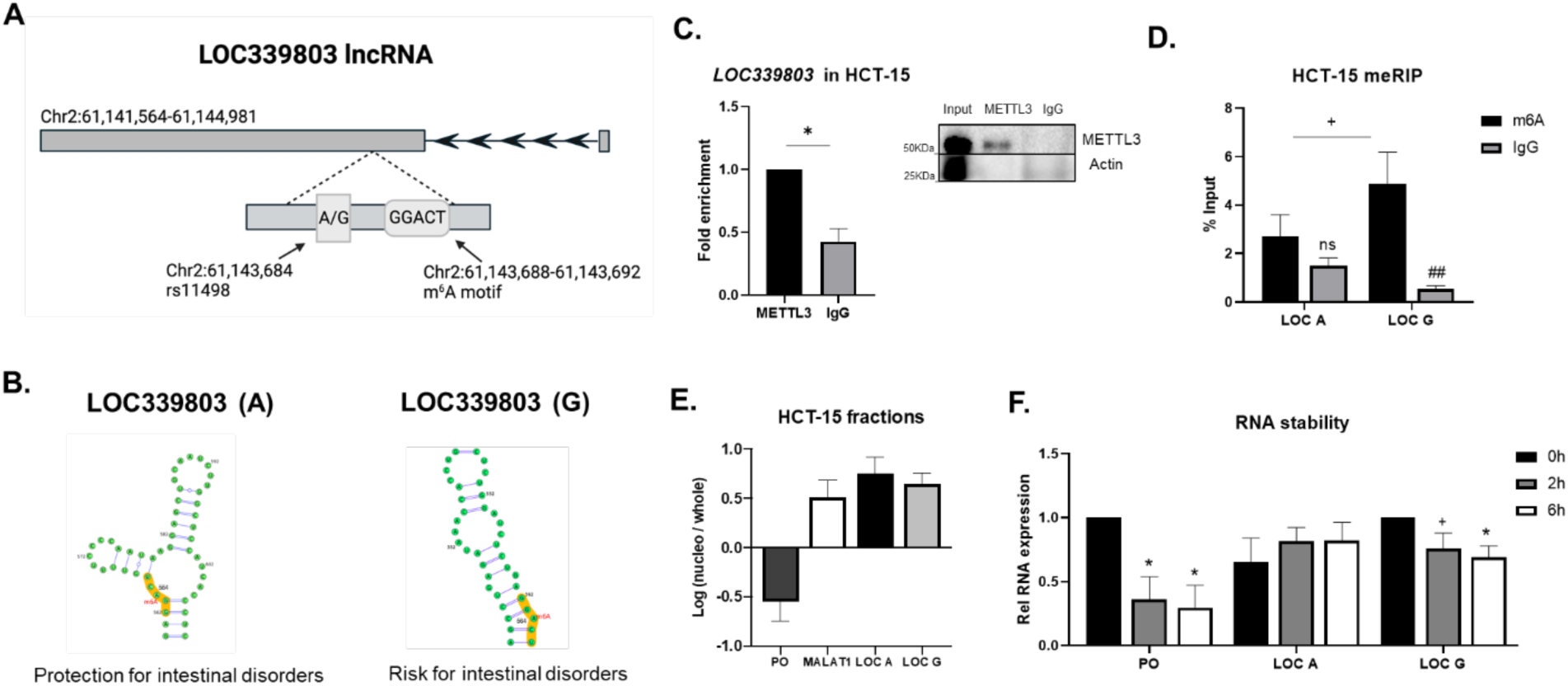
IBD-associated rs11498 SNP genotype affects m^6^A methylation levels in *LOC339803*. (**A**) Graphical representation of *LOC339803* chromosomic location created with BioRender. Immunochip SNP rs11498 and the GGACT m^6^A motif next to the SNP are zoomed in. **(B)** Predicted allele-specific secondary structure of the highly methylated motif region in *LOC339803* according to m^6^A predictor SRAMP online tool. **(C)** METTL3 RNA immunoprecipitation (RIP) and quantification of bound *LOC339803* levels assessed by RT-qPCR in HCT-15 intestinal cells. Right, representative immunoblot of the RIP experiments with Actin as negative control for the RIP. Data are means ± SEM (n=3 independent experiments). p-value determined by one-tailed Student’s t-test. **(D)** Allele-specific m^6^A methylation levels of *LOC339803* assessed by RNA immunoprecipitation using anti-m^6^A antibody in HCT-15 intestinal cells. Data are means ± SEM (n=4 independent experiments). p-value determined by two-way ANOVA test. **(E)** Subcellular localization of both *LOC339803* forms (*LOC A* and *LOC G*) using *P0* (*RPLP0,* cytoplasmic) and *MALAT1* (nuclear) as controls in HCT-15 intestinal cells. Data are means ± SEM (n=3 independent experiments). **(F)** RNA stability assay in HCT-15 cells treated with actinomycin for 2h and 6h using *RPLP0* as a positive control. Data are means ± SEM (n=3 independent experiments). p-value determined by one-tailed Student’s t-test. +p<0.1 *p<0.05; Enrichment relative to control IgG ##p<0.01.

Apart from differences in expression abundance, lncRNA localization also plays a critical role in their function and m^6^A methylation can influence methylated RNA localization (*29*). Subcellular localization assessment showed that both allelic forms of *LOC339803* are mainly nuclear in HCT-15 intestinal cells (Fig. 1E). When the stability of the lncRNA was studied, we could observe that while *LOC A* was quite stable, the preferentially methylated form *LOC G* significantly decreased upon incubation with actinomycin (Fig. 1F); suggesting increased methylation could lead to reduced stability of this nuclear lncRNA (*30*).

Hence, we confirmed that the IBD-associated rs11498 SNP genotype influences m^6^A methylation levels on *LOC339803* and that this differential methylation seems to influence the stability of the nuclear *LOC339803*.

### YTHDC1 m^6^A reader interacts with *LOC339803* influencing its cellular localization and protein binding

It is widely known that m^6^A methylated RNAs are recognized and bound by the so-called m^6^A readers to influence their function. The nuclear YTHDC1 m^6^A reader protein has been described to regulate a wide variety of RNA processes. More specifically, binding to the C-terminus of YTHDC1 has been proved to affect mRNA export (*29*), while interaction with the N-terminus of YTHDC1 has been linked with splicing or *XIST* mediated gene repression (*31, 32*). Given that YTHDC1 influences chromatin associated RNA stability (*30*) as well as the binding to target chromatin sites (*30, 31, 33*), we wondered whether this nuclear reader could be interacting with *LOC339803* in our intestinal model (Fig. S2A). YTHDC1 RIP confirmed its interaction with *LOC339803* (Fig. S2B), showing preferentially binding to the G allele (Fig. 2A), concordant with our previous observations (Fig. 1D). Previous observations by Roundtree et al. (*29*), described that binding to N-terminus of YTHDC1 could retain methylated RNAs in the nucleus. Thus, we further studied the effect of YTHDC1 specific binding in the subcellular localization of *LOC339803*. We observed that overexpression of the C-terminus in HCT-15 cell line resulted in a redistribution of our nuclear *LOC339803* towards the cytoplasm (Fig. 2B), supporting its interaction with the N-terminus and suggesting that differential binding of *LOC339803* to YTHDC1 termini could influence its localization, what could in turn regulate its function.

**Fig. 2.**
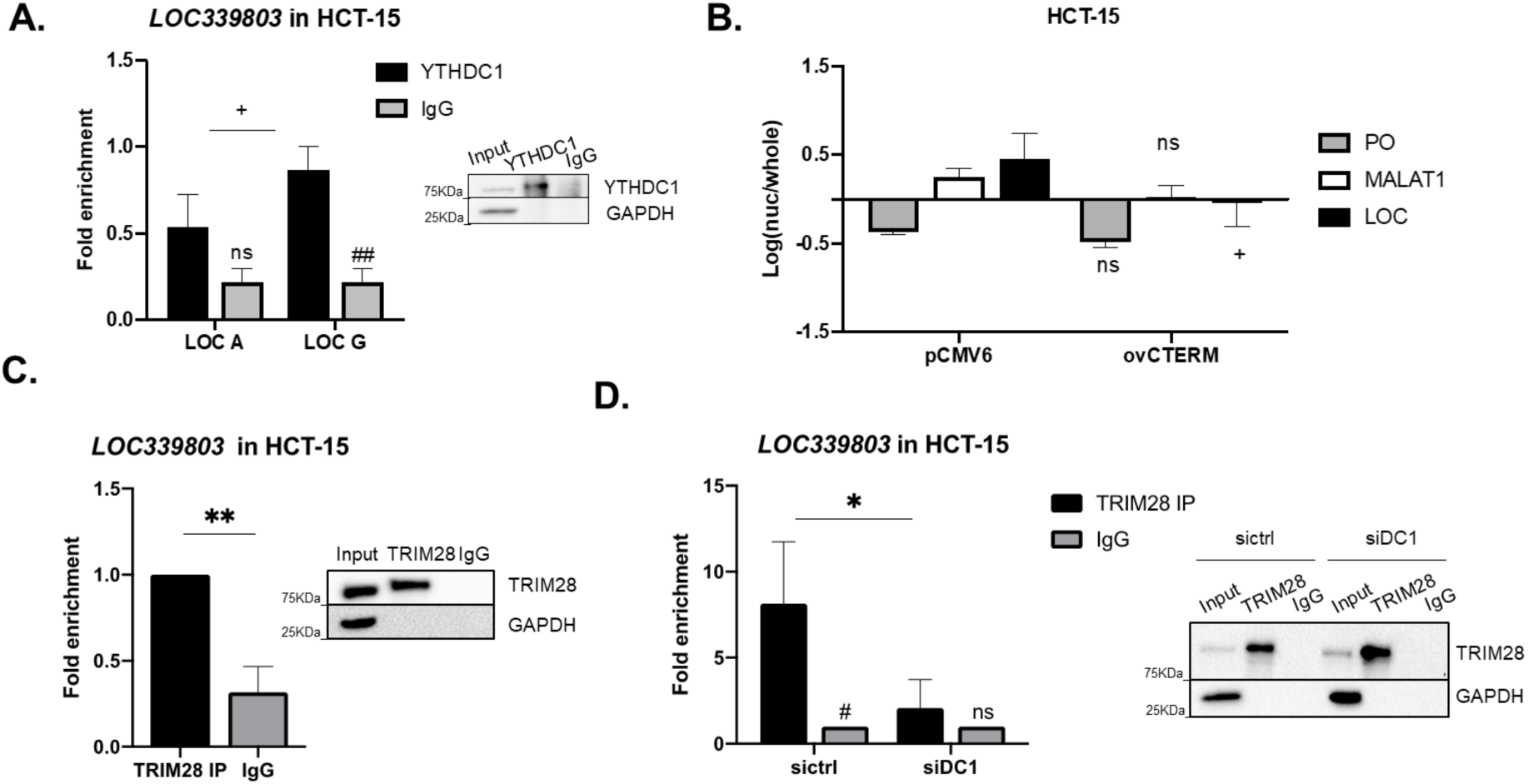
YTHDC1 m^6^A reader influences cellular localization and protein binding of *LOC339803*. **(A)** YTHDC1 RIP and quantification of allele-specific *LOC339803* levels assessed by RT-qPCR in HCT-15 intestinal cells. Right, representative immunoblot of the RIP experiment with GAPDH as negative control for the IP. Data are means ± SEM (n=4 independent experiments). p-values determined by two-way ANOVA test. **(B)** Subcellular localization of *LOC339803* using *P0* (*RPLP0,* cytoplasmic) and *MALAT1* (nuclear) as controls upon YTHDC1 C-terminus overexpression in HCT-15 intestinal cells. Data are means ± SEM (n=4 independent experiments). p-values determined by two-way ANOVA test. **(C)** TRIM28 RIP and quantification of bound *LOC339803* levels assessed by RT-qPCR in HCT-15 intestinal cells. Right, representative immunoblot of the RIP experiment with GAPDH as negative control for the IP. Data are means ± SEM (n=4 independent experiments). p-values determined by one-tailed Student’ t-test. **(D)** TRIM28 immunoprecipitation and quantification of bound *LOC339803* levels assessed by RT-qPCR in control (sictrl) or *YTHDC1* silenced (siDC1) HCT-15 cells. Right, representative immunoblot of the RIP experiment with GAPDH as negative control for the IP. Data are means ± SEM (n=4 independent experiments). p-values determined by two-way ANOVA test. +p<0.1, *p<0.05, **p<0.01; Enrichment relative to control IgG #p<0.05, ##p<0.01.

Given that YTHDC1 has been linked to interact with nuclear lncRNAs and affect their transcriptional regulatory roles (*30, 31, 34*), we wanted to further study the effect of YTHDC1 and *LOC339803* interaction. Pulldown of proteins interacting with *LOC339803* in HCT-15 cell line followed by mass spectrometry confirmed that *LOC339803* interacts with a wide variety of proteins (Fig. S2C, Sup Table 1). Moreover, Gene Ontology (GO) term analysis (*35–37*) showed an enrichment in nucleic acid binding, transcription machinery binding or chromatin and histone binding (Fig. S2C), suggesting *LOC339803* could participate in transcriptional regulation. Interestingly, pulldown-MS results showed that nuclear *LOC339803* binds TRIM28 and HDAC1 transcriptional repressors in HCT-15 cells, binding that was further confirmed by RIP experiments (Fig. 2C, S2D). These proteins had been previously identified to interact with YTHDC1 and to bind methylated RNAs (*34*). Indeed, silencing of YTHDC1 in HCT-15 cells (Fig. S2E) lead to a reduced interaction of *LOC339803* and TRIM28 (Fig. 2D), indicating that YTHDC1 is essential for *LOC339803* interaction with transcription regulators in intestinal cells.

Altogether, these results demonstrate that allele-specific differential methylation of *LOC339803* influences YTHDC1 binding ability. Additionally, we also confirmed that YTHDC1 is necessary for the binding of *LOC339803* to transcription regulators in intestinal cells, emphasizing the key role of m^6^A in the function of the lncRNA.

### *LOC339803* induction promotes transcriptional repression of *COMMD1* activating NFκB proinflammatory pathway

Having confirmed that *LOC339803* can bind transcription repressor proteins by a m^6^A dependent mechanism, we further studied *LOC339803* mechanism in intestinal cells. Assessment of its localization revealed that being primary nuclear, as we had already observed, it was equally distributed between the nucleoplasm and the chromatin (Fig 3A). Nuclear lncRNAs generally act in cis, regulating the transcription of their neighboring genes, and we had already observed that it binds transcription repressor proteins (Fig. 2C, S2C, S2D). Therefore, we analyzed the expression of intestinal inflammation related genes located nearby *LOC339803* after overexpression of the lncRNA (Fig. S3A). Among the analyzed genes, only *COMMD1* presented significant expression changes in response to *LOC339803* overexpression (Fig. 3B, S3B), with decreased levels that were more pronounced in the presence of the risk allele (*G).

**Fig. 3.**
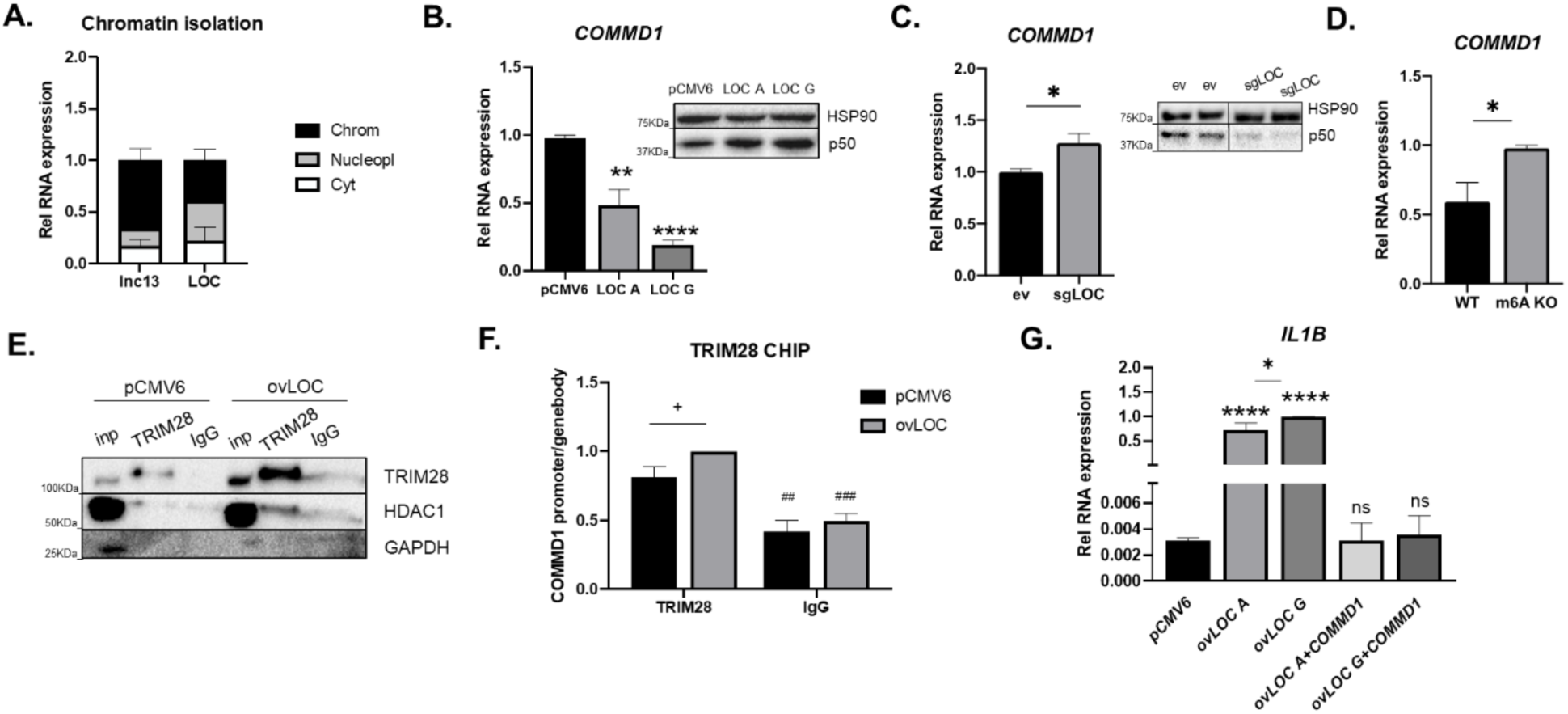
*LOC339803* induction promotes transcriptional repression of *COMMD1* activating NFkB proinflammatory pathway in intestines. (**A**) Subcellular localization of *LOC339803* using *lnc13* lncRNA (chromatin associated) as control in cytoplasmic, nucleoplasm and chromatin fractions from HCT-15 intestinal cells. Data are means ± SEM (n=3 independent experiments). **(B)** Left, quantification of *COMMD1* RNA levels by RT-qPCR and right, representative immunoblot of NFkB subunit p50 protein levels in cells transfected with overexpression plasmids of both forms of *LOC339803* (*LOC A* and *LOC G*). HSP90 was used as loading control. Data are means ± SEM (n=6 independent experiments). p-values determined by two-tailed Student’s t-test. **(C)** Left, quantification of *COMMD1* RNA levels by RT-qPCR and right, representative immunoblot of NFkB subunit p50 protein levels in cells with depleted *LOC339803* using CRISPR-Cas9. HSP90 was used as loading control. Data are means ± SEM (n≥5 independent experiments). p-value determined by one-tailed Student’s t-test. **(D)** Quantification of *COMMD1* RNA levels by RT-qPCR in cells with a deletion of the m^6^A methylation region in *LOC339803* using CRISPR-Cas9. Data are means ± SEM (n=4 independent experiments). p value determined by two-tailed Student’s t-test. **(E)** Representative immunoblot of the co-immunoprecipitation of TRIM28 and HDAC1 using anti-TRIM28 antibody in cells transfected with an empty vector (pCMV6) or *LOC339803* overexpressing plasmids (ovLOC). GAPDH was used as a negative control for the CO-IP. **(F)** Chromatin immunoprecipitation (ChIP) using anti-TRIM28 antibody and quantification of bound *COMMD1* promoter levels in cells transfected with an empty vector (pCMV6) or *LOC339803* overexpressing plasmids (ovLOC). Data are means ± SEM (n=3 independent experiments). p value determined by two-way ANOVA test. **(G)** Quantification of *IL1B* RNA levels by RT-qPCR in cells transfected with overexpression plasmids of both forms of *LOC339803* (*LOC A* and *LOC G*) and *COMMD1* (*ovLOC A* + *COMMD1* and *ovLOC G* + *COMMD1*). Data are means ± SEM (n=3 independent experiments). p values determined by one-way ANOVA test. +p<0.1, *p<0.05, **p<0.01, ****p<0.0001. Enrichment relative to control IgG ##p<0.01, ###p<0.001.

Interestingly, *COMMD1* repression is known to be related with NFκB activation (*38*), which is a central player in the development of inflammatory diseases of the intestine (*39, 40*). In line with this, we observed augmented amounts of NFκB p50 subunit in the cells overexpressing *LOC339803,* with a significant increase only in the presence of the risk allele (*G) (Fig 3B, S3C). On the other end, cells with a total deletion of the lncRNA showed increased *COMMD1* expression together with a decrease in p50 levels (Fig 3C, S3D). Accordingly, cells with a CRISPR-Cas9 mediated deletion of the m^6^A motif in *LOC339803* presented induced *COMMD1* expression levels (Fig 3D, S3E), confirming the implication of *LOC339803* and the relevance of m^6^A methylation in the regulation of downstream inflammatory processes.

As these results pointed to a lncRNA-mediated transcriptional regulation of *COMMD1,* we wanted to describe how *LOC339803* regulates *COMMD1* expression. We had observed that YTHDC1 is necessary for *LOC339803*-TRIM28 interaction (Fig. 2D), underlying the importance of m^6^A methylation on lncRNA function. Indeed, DNase I hypersensitivity assay in HCT-15 cells confirmed that when the risk *LOC G* form (presenting higher methylation levels) is overexpressed, the DNase I hypersensitivity site within *COMMD1* promoter is less accessible, suggesting that the lncRNA could be contributing to a lower *COMMD1* transcription (Fig. S3F). In addition, TRIM28, which has been described to interact with YTHDC1 and to bind methylated RNAs, works as a scaffold protein to recruit diverse repressor proteins as histone deacetylases or H3K9 methyltransferase SETDB1 (*33, 41*). Using CO-IP, we were able to confirm that HDAC1 protein (which was also found bound to *LOC339803*) (Fig. S2D) interacts with TRIM28 in HCT-15 cells (Fig. 3E). Moreover, overexpression of *LOC339803* revealed stronger HDAC1-TRIM28 interaction (Fig. 3E) as well as an increased binding of TRIM28 to *COMMD1* promoter as assessed by chromatin immunoprecipitation (ChIP) (Fig. 3F) which is concordant with an inflammatory environment.

Collectively, these results showed that *LOC339803* binds to transcription repressor proteins, forming a transcription repressor complex that regulates *COMMD1* and induces NFκB. Considering that *IL1B* proinflammatory cytokine is induced by NFκB and is a key driver of IBD, to test whether *LOC339803* mediated *COMMD1* regulation affects inflammation; we analyzed the expression of *IL1B* in this scenario. We observed that overexpression of *LOC339803* was able to induce *IL1B* expression, being significantly higher in the presence of risk allele (*G). This induction was reverted when *COMMD1* was overexpressed (Fig. 3G) and silencing of *COMMD1* significantly induced *IL1B* expression (Fig. S3G); confirming that the induction of *LOC339803* augments the expression of *IL1B* via *COMMD1* repression.

Overall, these results show that the increased methylation present in the risk allele (*G) enhances the interaction of YTHDC1 with the lncRNA, favoring the binding of *LOC339803*-TRIM28-HDAC1 repressor complex to *COMMD1* promoter, finally resulting in increased NFκB and *IL1B* proinflammatory cytokine expression.

### IBD patients present increased *LOC339803* expression, which correlates with higher risk to develop GI malignancies

As the *in vitro* results obtained in intestinal cells pointed to an implication of *LOC339803* in the pathogenesis of IBD by the induction of *IL1B*, we next wanted to confirm these results using human patient samples. For this aim, intestinal biopsies from controls and patients with the two IBD subtypes (ulcerative colitis and Crohn’s disease) were analyzed.

We first observed that IBD patients present altered expression of several m^6^A machinery members (Fig. S4A), pointing to m^6^A methylation as an important player in disease related regulatory pathways (*24, 25*). Specifically, we observed that the expression of YTHDC1 reader is significantly increased in IBD patients. According to our *in vitro* results, this could enhance the binding of *LOC339803* and the repressor complex to *COMMD1* promoter, inducing an *IL1B* mediated proinflammatory environment in these individuals. Indeed, IBD patients also present increased *LOC339803* and reduced *COMMD1* expression levels (Fig. 4 A,B). Moreover, we confirmed that IBD patients also present increased *IL1B* expression (Fig. 4C), as a sign of increased proinflammatory environment in their intestines. Additionally, we could also confirm that the increased *LOC339803* levels correlate with high *IL1B* expression in human intestinal samples (Fig. S4B). Further confirming the implication of the lncRNA in the proinflammatory boost of IBD, we found that individuals carrying the risk allele in the lncRNA (GG/AG genotypes) present significantly higher *IL1B* levels in the intestine (Fig. 4D). Altogether, our results in IBD patients confirm that the increased inflammation present in these individuals is, at least in part, mediated by *LOC339803*-induced *IL1B* levels.

**Fig. 4.**
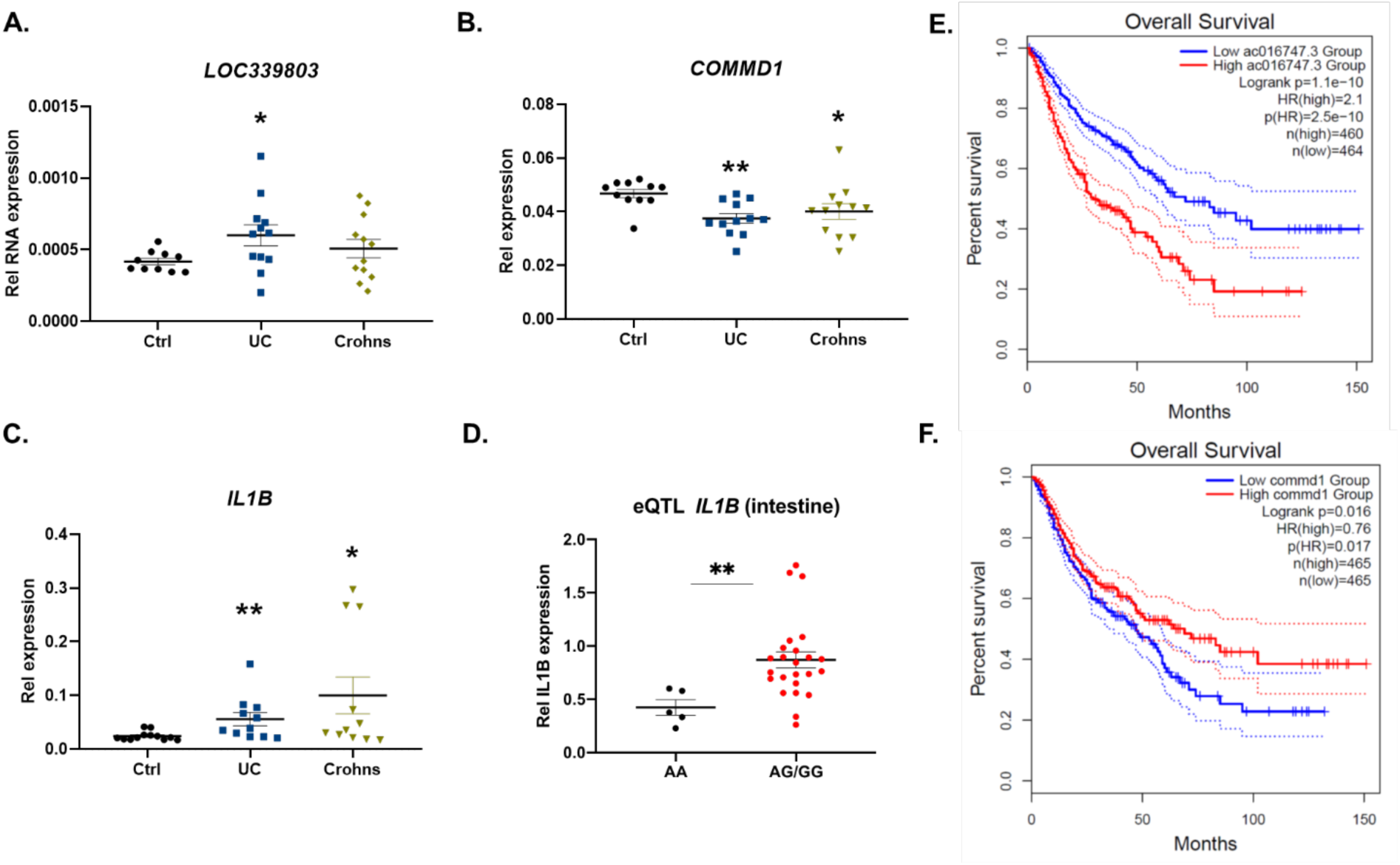
IBD patients present increased *LOC339803* expression which correlates with higher risk to develop GI malignancies. Quantification of **(A)** *LOC339803,* **(B)** *COMMD1* and **(C)** *IL1B* by RT-qPCR in intestinal biopsies from controls and patients with ulcerative colitis (UC) and Crohn’s disease (Crohns). Data are means ± SEM (n≥10). **(D)** eQTL of *IL1B* expression levels in intestinal samples comparing individuals with the protection genotype (AA) and individuals harboring the risk allele (AG+GG). Data are means ± SEM (n≥5). All p-values determined by Mann-Whitney test. *p<0.05, **p<0.01. **(E-F)** Overall survival graphs obtained from GEPIA (http://gepia.cancer-pku.cn/detail.php?gene=&clicktag=boxplot) for GI cancers (COAD=Colon adenocarcinoma; ESCA=Esophageal carcinoma; READ=Rectum adenocarcinoma; STAD=Stomach adenocarcinoma) for **(E)** *LOC339803* (*AC016747.3*) and **(F)** *COMMD1*.

It is known that chronic inflammation increases risk to develop other malignancies, as GI cancers in IBD patients. Interestingly, analysis of online data from GEPIA web app (*42*) showed that individuals with increased *LOC339803* (*AC016747.3*) levels have lower overall survival in GI cancers (Fig. 4E). Concordantly, low *COMMD1* levels also decrease the overall survival in these individuals (Fig. 4F). These results highlight the importance of a controlled regulation of the *LOC339803* induced inflammatory pathway to protect against the increased inflammation in IBD patients that could in turn derive in GI cancers. In accordance, and as observed in online data (*42*) most GI tumor tissues have increased *IL1B* (Fig. S4C).

To sum up, our results in human samples confirm the functional implication of *LOC339803* in IBD. Additionally, these results explain the association of the region harboring SNP rs11498 with IBD, pointing to *LOC339803* lncRNA as a key player in the inflammatory environment of this disease. Additionally, our data suggests that *LOC339803* expression levels could be involved in cancer related complications in IBD patients, emerging as an interesting therapeutic target for this disorder and is related comorbidities.

### *LOC339803* targeting as a therapeutic strategy

As the results in human samples confirmed the implication of *LOC339803* in the development of IBD, and considering the recent development on RNA based therapies, we wanted to analyze the putative therapeutic use of *LOC339803* and/or its methylation levels in an intestinal inflammatory disease model.

Results so far indicate that *LOC339803* induces *IL1B* proinflammatory cytokine and that m^6^A mediated interaction of the lncRNA with YTHDC1 reader protein is key for the proper function of *LOC339803*. Therefore, we aimed to address the role of m^6^A and *LOC339803* in the regulation of *IL1B* in an *in vitro* disease model.

We first confirmed that by manipulating m^6^A machinery (overexpression *METTL3*) or reducing total m^6^A levels using cycloleucine, *LOC339803* expression levels can be modulated (Fig. 5A,B). To verify the importance of m^6^A methylation in *LOC339803* proinflammatory functionality, we also built a *LOC339803* overexpressing plasmid with truncated m^6^A motif, so that the m^6^A mark next to the SNP would disappear (Fig. S5A). We confirmed the involvement of the m^6^A motif for the proper function of the lncRNA in intestinal cells, as the overexpression of the m^6^A mutant plasmid did not induce *IL1B* levels as it was observed with the wild type plasmid (Fig 5C, S5B). These results confirm the importance of m^6^A methylation in *LOC339803* functionality as well as its significance as a potential therapeutic target and highlight m^6^A-targeted drugs as a good approach to regulate *LOC339803* levels in intestinal inflammatory disorders as IBD.

**Fig. 5.**
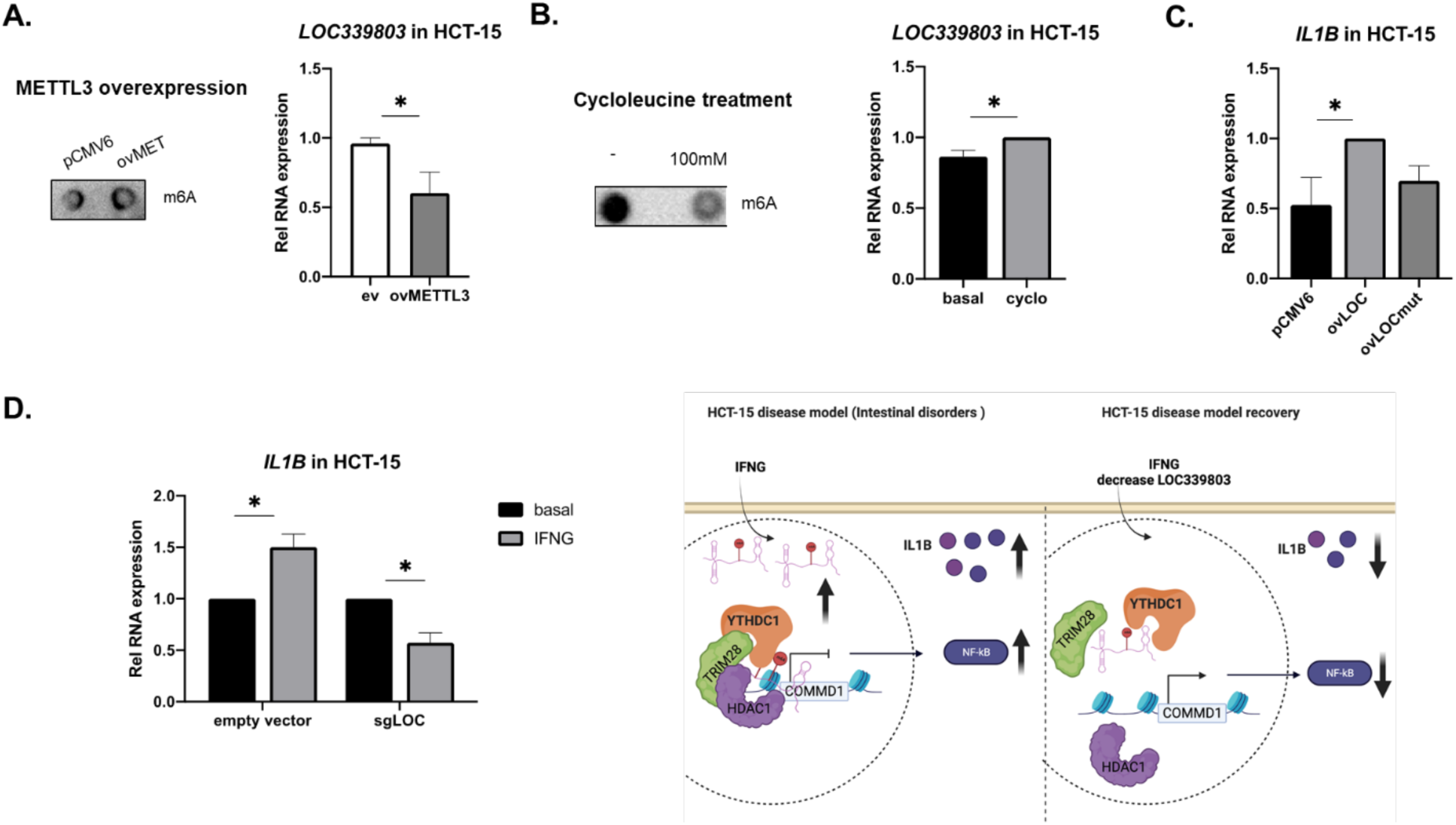
*LOC339803* targeting as a therapeutic strategy. **(A)** Left, quantification of total m^6^A levels upon METTL3 overexpression by dot-blot. Right, quantification of *LOC339803* RNA levels by RT-qPCR in METTL3 overexpressing cells (ovMETTL3) in HCT-15 cells. Data are means ± SEM (n=5 independent experiments). p-value determined by one-tailed Student’s t-test. **(B)** Left, quantification of total m^6^A levels after cycloleucine treatment by dot-blot. Right, quantification of *LOC339803* RNA levels by RT-qPCR in basal or cycloleucine treated cells in HCT-15 cells. Data are means ± SEM (n=3 independent experiments). p-value determined by one-tailed Student’ t-test. **(C)** Quantification of *IL1B* RNA levels by RT-qPCR in cells transfected with an empty vector (pCMV6), *LOC339803* overexpressing plasmid (ovLOC) or *LOC339803* overexpressing plasmid with mutated m^6^A motif (ovLOCmut) in HCT-15 cells. Data are means ± SEM (n=4 independent experiments). p-values determined by one-way ANOVA test. (**D**) *In vitro* stimulation model in HCT-15 intestinal cells using interferon gamma (IFNG). Left, quantification of *IL1B* RNA levels by RT-qPCR in cells transfected with an empty vector or *LOC339803* depleted cells using CRISPR-Cas9 (sgLOC). Data are means ± SEM (n=3 independent experiments). Right, representative image of stimulation and recovery models in HCT-15 cells created with Biorender. p-values determined by one-tailed Student’s t-test. *p<0.05.

To further assess the effect of *LOC339803* in the activation of a proinflammatory response in IBD, we used an interferon gamma (IFNG) stimulation model, described to alter m^6^A methylation levels in intestinal cells (*25*). With the aim of reducing IFNG-induced inflammation in these cells, we knocked down *LOC339803* by CRISPR-Cas9 in HCT-15 cells (Fig. S5C). We observed that *LOC339803* decrease prevented the induction of the proinflammatory cytokine *IL1B* upon IFNG stimulation (Fig. 5D), confirming the participation of this lncRNA on *IL1B* induction in the intestine.

These results show that *LOC339803* and its m^6^A methylation levels are key to induce *IL1B* expression in intestinal epithelial cells and explain how this disease-specific risk variant contributes to the predisposition to disease development by generating higher basal *IL1B* levels. In addition, *LOC339803* evolves as an interesting therapeutic target for IBD, as modifying m^6^A levels and manipulating lncRNA expression seem to help prevent the pro-inflammatory environment that would lead to disease development.

## DISCUSSION

In this study, we have described that m^6^A methylation provides *LOC339803* with the ability to regulate the expression of the key proinflammatory *IL1B* cytokine via its interaction with YTHDC1 and transcription repressor proteins (Fig. S6). In addition, we have confirmed that *LOC339803* expression levels are deregulated in patients with IBD, and we have observed that the genotype of rs11498 SNP influences the expression levels of the downstream proinflammatory cytokine *IL1B*, predisposing individuals to chronic inflammation.

We have also confirmed that the genotype of the rs11498 SNP influences m^6^A methylation levels on *LOC339803* and that this lncRNA shows a methylation-dependent function. RNA modifications are emerging as a new regulatory layer, playing critical roles in RNA processing, splicing, translation or stability (*43, 44*), all of them crucial in the development of multiple human diseases (*45, 46*). m^6^A-QTLs are known to contribute to the risk to develop some immune and blood-related disorders; being enriched within RNA-protein binding sites, RNA structure-changing variants and transcriptional features (*17*). In addition, SNPs have been shown to be enriched in lncRNAs and studies have proved that associated SNPs can alter lncRNA functions, most probably by altering secondary structures and/or their binding ability (*12, 13, 47*). Indeed, in accordance with previous results, we have observed that the more methylated *LOC G* form is less stable, most probably due to increased interaction with YTHDC1 reader described to alter the stability of m^6^A methylated RNAs (*30*). These results show how the genotype dependent m^6^A levels can alter the lncRNA function, as we described that allele-specific differential methylation of *LOC339803* determines YTHDC1 binding ability and this interaction is necessary for the coupling of *LOC339803* to the transcription repressor complex. Based on our results and considering that m^6^A-QTLs seem to affect key aspects of lncRNA functionality, identification of disease associated SNPs within m^6^A methylation motifs seem to be a good approach to identify crucial players in key pathological pathways.

Cell type specific mechanisms to activate *IL1B* have been described in monocytes and macrophages and the different cellular environment has been proposed as the reason of this regulatory setup (*48*). Our results indicate that *LOC339803* is, at least in part, responsible for the regulation of *IL1B* in intestinal cells. While the role of immune cells in the innate immune response has been known for a long time, increasing evidence indicate the importance of endothelial and epithelial cells in the early steps of inflammatory response (*49*). Hence, production of cytokines by epithelial cells and their role in early response steps are important to clarify the development of inflammatory disorders. In this work, we have shown that intestinal epithelial cells express *IL1B* proinflammatory cytokine, what will increase the proinflammatory environment and attract immune cells, increasing the risk to develop other GI malignancies. In addition, we observed that the risk genotype leads to increased *IL1B* production predisposing the individuals to augmented basal intestinal inflammation that will contribute to the predisposition to IBD. We also proved the transcriptional regulatory function of the nuclear *LOC339803* in intestinal cells. *LOC339803* forms a transcription repressor complex with TRIM28 and HDAC1 to reduce *COMMD1* levels, leading to higher levels of the proinflammatory cytokine *IL1B*, characteristic of intestinal inflammation (*50*). Whether this mechanism can also regulate other inflammatory genes could be an interesting follow-up study.

Lately, the interest on RNA based therapies, mainly those based on small RNA and mRNA vaccines is increasing. Considering the abundance of lncRNAs and their involvement in cellular biology and human diseases, lncRNAs are noteworthy therapeutic targets. Their low expression profiles and tissue-specificity make them appealing, as low dosages should be enough for disease treatment (*51*). This study highlights the putative therapeutic use of *LOC339803* in IBD. Reduction of the lncRNA helped lowering the IFNG induced inflammatory response in intestinal cells. Therefore, the use of small molecules that disrupt lncRNA function or antisense oligonucleotides that lower their expression seem good tools for lncRNA focused therapies. We also showed that deletion of the methylation motif in *LOC339803* favored a lesser inflammatory response in intestinal cells. Hence, m^6^A methylation as well as the nearby associated SNP region seem key for proper lncRNA function. The different clinical trials based on lncRNA therapies already being carried out, strengthen our results as a starting point for the development of the so needed therapies for IBD treatment. Moreover, we also present m^6^A targeted approaches as another interesting alternative therapy as many m^6^A-targeted drugs are already being tested (*52*).

One of the main drawbacks of our work, that also applies to many others studying lncRNAs (*53*), is the lack of a murine homolog for *LOC339803*, making it implausible to perform preclinical studies in murine disease models. Nevertheless, we believe that the studies performed using human *in vitro* models and the confirmation in human intestinal samples make our model strong and present enough evidence to conclude that *LOC339803* is involved in disease pathogenesis and could be a good therapeutic target. To sum up, our results show the implication of lncRNAs harboring disease associated variants and epitranscriptomic modifications, such as m^6^A RNA methylation, to respond to external stimuli and to activate important inflammatory response players, as *IL1B* cytokine. These results highlight the importance of studies in which the functional implication of a lncRNA is described in disease scenario and emphasize the importance of lncRNA and m^6^A methylation function in IBD pathogenesis opening the door to novel therapeutic alternatives.

## MATERIALS AND METHODS

### Study design

In this study, human bowel biopsy samples from inflammatory bowel disease and control individuals were obtained from Hospital de Galdakao-Usansolo (Spain). Experiments using these biopsy samples were approved by the corresponding Ethical Committee. For *in vitro* experiments human intestinal cell line HCT-15 (Sigma-Aldrich, # 91030712) was used. Disease model stimulations were performed using IFNG in HCT-15. All the *in vitro* experiments were performed at least three independent times.

### m^6^A RNA immunoprecipitation

4 μg of precleared RNA per sample were fragmented with RNA fragmentation buffer (100 mM Tris, 2 mM MgCl_2_) for 3 min at 95°C and placed on ice immediately after heating. 10% of RNA was kept as input. 1 µg of m^6^A antibody (Abcam, #ab151230) and control antibody (IgG, Santa Cruz Biotechnologies, Dallas, USA, #sc-2025) were coupled to agarose A beads (GE Healthcare, Chicago, USA) in a rotation wheel for 1 h at 4°C. After incubation, beads were washed twice in reaction buffer (150 mM NaCl, 10 mM Tris-HCl, 0.1 % NP-40). RNA was added to the antibody-coupled beads and incubated for 3 h at 4°C in a rotating wheel. Subsequently, beads were washed 3X in reaction buffer, 3X in low salt buffer (50 mM NaCl, 10 mM TrisHCl and 0.1 % NP-40) and 3X in high salt buffer (500 mM NaCl, 10 mM TrisHCl and 0.1 % NP-40). After the last wash, beads were resuspended in Lysis buffer and RNA was extracted using the PureLink RNA extraction kit (Invitrogen, Carlsbad, USA, #12183016).

### RNA immunoprecipitation assay (RIP)

For RIP experiments, HCT-15 cells were lysed in RIP buffer (150 mM KCl, 25 mM Tris, 0.5 mM DTT, 0.5 % NP-40, PI), kept on ice for 15 minutes and homogenized using a syringe. Lysates were pre-cleared with protein A-Agarose beads (GE Healthcare, Chicago, USA) for 1 h in a wheel shaker at 4°C. A-Agarose beads were blocked with 20 % BSA and mixed with pre-cleared lysates and 1 µg of anti-IgG antibody (negative control; Santa Cruz Biotechnologies, #sc-2025) or antibody of interest. After overnight incubation in a wheel shaker at 4°C, beads were washed 3X with RIP buffer, 3X with low salt buffer (50 mM NaCl, 10 mM Tris-HCl, 0.1 % NP-40) and 3X with high salt buffer (500 mM NaCl, 10 mM Tris-HCl, 0.1 % NP-40). After the washes, 70 % of beads were resuspended in RNA extraction buffer and 30 % was used for WB.

### Chromatin immunoprecipitation assay (ChIP)

For ChIP experiments, HCT-15 cells were crosslinked with formaldehyde and collected in PBS with a scratcher. Cell pellet was then resuspended in L1 buffer (50 mM Tris pH8, 2 mM EDTA, 0.1 % NP-40, 10 % glycerol) + PI and incubated in ice for 5 min. Supernatant was discarded and pellet resuspended in 300 uL L2 Buffer (50 mM Tris pH8, 0.1 % SDS, 5 mM EDTA) + PI to disrupt the chromatin using bioruptor sonicator. Centrifuged samples at maximum speed were used for immunoprecipitation.

ChIP dilution buffer (50 mM Tris pH8, 0.5 % NP-40, 0.2 M NaCl, 0.5 mM EDTA) was added up to 1 mL and in order to reduce non-specific background, the samples were pre-incubated with 60 uL of protein A-Agarose beads (GE Healthcare, Chicago, USA) + Salmon Sperm DNA (Invitrogen #15632-011) (1 ug of DNA/20 uL of protA) for 60 mins at 4°C shaking. The supernatant was collected and equal volumes were put into 2 tubes.

A negative control antibody (IgG, Santa Cruz Biotechnologies, #sc-2025) or the antibody of interest (TRIM28, Santa Cruz Biotechnologies, #sc-515790) and 60 uL of blocked protein A-Agarose beads (with 10% BSA) were added to lysate and incubated overnight at 4°C on a shaker. After overnight incubation, beads were washed 3X with high salt wash buffer (20 mM Tris pH8, 0.1 % SDS, 1 % NP-40, 2 mM EDTA, 0.5 M NaCl) and 3X with TE buffer (10X TE buffer: 0.1 M Tris HCl, 0.001 M EDTA pH8). After the washes, the beads were resuspended in NTI buffer for DNA purification using NucleoSpin Gel and PCR Clean-up kit (Macherey-Nagel, #740609.250).

### *LOC339803* KO cell generation using CRISPR Cas9

For *LOC339803* KO cell line generation two sgRNAs flanking the lncRNA were designed and cloned in a px459 vector. HCT-15 cells were transfected with 250 ng of each vector and selected with puromycin. After selection, clonal cell lines were generated by serial dilution. The sequences for the sgRNAs are shown in Sup Table 2.

### m^6^A KO cell generation using CRISPR Cas9

For m^6^A KO cell line generation, two sgRNAs flanking the m^6^A motif were designed and cloned in px458 GFP and px330 mCherry vectors. HCT-15 cells were transfected with 250 ng of each plasmid. HCT-15 cells were sorted by cell sorter BD FACSJazz (2B/4YG) 48 hours post-transfection for the generation of clonal cell lines. The sequences for the sgRNAs are shown in Sup Table 2.

### Statistical analysis

All the statistical analyses were performed using GraphPad Prism 8 (GraphPad Software). Significance was calculated using Student’s t-test, Mann Whitney test or ANOVA test as specified in figure legends. The statistically significance level was set at p< 0.05. p-values lower than 0.1 are marked with a +.

## SUPPLEMENTARY MATERIALS

### MATERIALS

#### Cell lines and treatments

Intestinal HCT-15 (#91030712) cell line was purchased from Sigma-Aldrich (Poole, UK) and cultured in RPMI (Lonza, #12-115F) with 10 % FBS, 100 units/ml penicillin and 100 μg/ml streptomycin.

For the intestinal inflammation model, WT and CRISPR Cas9 mediated *LOC339803* deleted HCT-15 cells were stimulated with 100 U of IFNG for 30 min.

#### Human patients and samples

The patients and the public were engaged during recruitment of volunteers via informative flyers to patients and at outreach activities open to the general public. For intestinal samples, participation in the study was offered to patients scheduled to a routine gastroscopy/colonoscopy with symptomatic indication in the Endoscopy Unit of Galdakao University Hospital. The findings of the study will be disseminated in events organized by the societies and in National and International meetings.

For IBD patients and controls, individuals in high risk of undiagnosed or already diagnosed IBD patients, extra biopsy specimen was obtained in routine colonoscopy of the inflamed segment if present. None of the patients suffered from any other concomitant immunological disease. None of the controls showed small intestinal inflammation at the time of the biopsy.

This study was approved by the Basque Country Clinical Research Ethics Board (CEIC-E ref. PI2019133) and analyses were performed after informed consent was obtained from all subjects. All experiments were performed in accordance with relevant guidelines and regulations.

### METHODS

#### DNA, RNA and protein extraction

For human biopsies NucleoSpin TriPrep kit (Macherey-Nagel, Düren, Germany, #740966.50) was used following manufacturer instructions. For HCT-15 RNA extraction was performed using NucleoSpin RNA Kit (Macherey Nagel, #740984.50) and cells were lysed in RIPA buffer for protein quantification.

#### Gene expression analyses

500-1000 ng of RNA were used for the retrotranscription reaction using iScript cDNA Synthesis Kit (BioRad, CA, USA, #1708890). Expression values were determined by RT-qPCR using Sybr Green (iTaq SYBR Green Supermix, Bio-Rad, #1725124) and specific primers. *RPLP0* gene was used as endogenous control in both human samples and cell lines. Reactions were run in a BioRad CFX384 and melting curves were analyzed to ensure the amplification of a single product. All qPCR measurements were performed in duplicates and expression levels were analyzed using the 2^−ΔΔCt^ method and normalization to the highest value was used for relative RNA expression calculation. All primers are listed in Sup Table 2.

#### SNP genotyping

Genotyping of the SNP rs11498 was performed in DNA samples of human biopsies using a custom Taqman SNP Genotyping Assay (ThermoFisher, Waltham, MA) following the manufacturer’s instructions.

#### Western Blot

Laemmli buffer (62 mM Tris-HCl, 100 mM dithiothreitol (DTT), 10 % glycerol, 2 % SDS, 0.2 mg/ml bromophenol blue, 5 % 2-mercaptoethanol) was added to the protein extracts in RIPA and were denatured by heat. Proteins were migrated on 10% SDS-PAGE gels. Following electrophoresis, proteins were transferred onto nitrocellulose membranes using a Transblot-Turbo Transfer System (Biorad) and blocked in 5 % non-fatty milk diluted in TBST (20 mM Tris, 150 mM NaCl and 0.1 % Tween 20) at room temperature for 1 h. The membranes were incubated overnight at 4°C with primary antibodies diluted 1:1000 in TBST. Immunoreactive bands were revealed using the Clarity Max ECL Substrate (BioRad, #1705062) after incubation with a horseradish peroxidase-conjugated anti-mouse or anti-rabbit (1:10000 dilution in 2.5 % non-fatty milk) secondary antibody for 1 h at room temperature. The immunoreactive bands were detected using a Bio-Rad Molecular Imager ChemiDoc XRS and quantified using the ImageJ software. The following antibodies were used for Western Blotting: YTHDC1 (Abcam, #264375, Cell Signalling, #E4I9E), HSP90 (Cell Signaling; #4874), GAPDH (Santa Cruz Biotechnologies, #sc-47724), TRIM28 (Santa Cruz Biotechnologies, #sc-515790), HDAC1 (proteintech, #66085-1), p50 (Santa Cruz Biotechnologies, #sc-8414), METTL3 (Abcam, #195352), ACTIN (Santa Cruz Biotechnologies, #sc47778).

#### Plasmid construction and overexpression

*LOC339803* was amplified from human cDNAs containing A or G allele for rs11498 SNP and cloned into a pCMV6 vector (Origene, #PS100001) using KpnI and FseI restriction sites. For *LOC339803* overexpression plasmids with the truncated m^6^A motif, the commercially available Phusion^TM^ Site-Directed Mutagenesis Kit (#F541) was used. For YTHDC1 C-term construct the YTH domain together with the C-terminal were cloned in a CMV driven vector using AscI and FseI restriction sites. The primers used for cloning are listed in Sup Table 2.

For C-termini overexpression experiments 400 ng plasmid were used. 300.000 cells/well were seeded and transfected using X-TremeGENE HP DNA transfection reagent (Sigma-Aldrich, #6366546001), cells were harvested after 48 h.

For *LOC339803* and mutant *LOC339803* overexpression, 200 ng of plasmids per 100.000 cells were used. Cells were seeded and transfection was performed with X-TremeGENE HP DNA transfection reagent (Sigma-Aldrich, #6366546001) for 24 or 48 h.

For *COMMD1* overexpression experiments, 250 ng plasmid from Origene (#RC205614) was used. 150.000 cells/well were seeded and transfected using X-TremeGENE HP DNA transfection reagent (Sigma-Aldrich, #6366546001), cells were harvested after 24 h.

#### Silencing experiments

For *YTHDC1*, *LOC339803* or *COMDD1* silencing, 30 nM of 2 different siRNAs against *YTHDC1* (IDT, #hs.Ri.YTHDC1.13.1 and hs.Ri.YTHDC1.13.3), *LOC339803* (IDT, #hs.Ri.LOC339803.13.1 and hs.Ri.LOC339803.13.2), *COMMD1* (IDT, #hs.Ri.COMMD1.13.2 and hs.Ri.COMMD1.13.3) or negative control siRNA (IDT #51-01-14-01) were transfected into cells using Lipofectamine RNAimax reagent (Invitrogen).

#### Co-immunoprecipitation assay (CO-IP)

For CO-IP experiments, HCT-15 cells were lysed in CO-IP buffer (20 mM Tris pH8, 150 mM NaCl, 1mM EDTA, 1mM EGTA, 1% Triton X-100) and the same protocol described for RIP was followed. TRIM28 (Santa Cruz Biotechnologies, #sc-515790) antibody was used for CO-IPs.

#### RNA immunoprecipitation assay followed with mass spectrometry (RIP-MS)

For RIP-MS experiments, sense and antisense *LOC339803* were amplified from cDNA using a T7 promoter primer. The PCR product was purified and used for in vitro transcribing biotinylated RNA using the T7 polymerase (Takara) and RNA biotin labeling kit (Roche). 1 μg of purified *LOC339803* RNA was mixed and incubated with cell extracts from HCT-15 cell. Streptavidin beads were added to the reaction and further incubated. After incubation, beads were washed 5 times. Samples were sent for mass spectrometry and subjected to in-solution digestion followed by nano LC-MS/MS analysis. List of RIP-MS results are in sup. Table 1.

#### Cellular fractionation

For the quantification of RNA amounts in nuclear and cytoplasmic compartments, nuclei were isolated using C1 lysis buffer (1.28 M sucrose, 40 mM Tris-HCl pH 7.5, 20 mM MgCl_2_, 4 % Triton X-100). The amounts of *LOC339803-A/G*, *MALAT1* (nuclear control) and *RPLP0* (cytoplasmic control) were measured by RT-qPCR and compared to the total amount of those RNAs in the whole cell lysate.

For the quantification of RNA amounts in chromatin and nucleoplasm compartments, HCT-15 cells were crosslinked using 16% formaldehyde. After crosslinking, cells were centrifuged and resuspended in NARA buffer (500 mM HEPES pH7.9, 1 M KCl, 500 mM EDTA, 0.05 % NP-40) for cytoplasm separation. Nuclei were resuspended in low-salt buffer (10 mM Tris HCl pH 7.4, 0.2 mM MgCl_2_, 1 % triton) and after centrifugation nucleoplasm was transferred to fresh tubes. Chromatin was resuspended in HCl 0.2 N centrifuged and neutralized with 1 M Tris-HCl pH8. Obtained lysates were decrosslink prior to RNA extraction.

For the quantification of protein amounts in nuclear and cytoplasmic compartments, cells were resuspended in NARA buffer (10 mM HEPES pH 7.9, 10 mM KCl, 0.1 mM EDTA) with PI and incubated in ice for 10 minutes. After adding NP-40 to final concentration 0.05 %, lysates were incubated 5 minutes in ice and centrifuged at 400 g for 2 minutes. The supernatant was the cytosolic fraction. Pellet was washed 3X with NARA buffer and resuspended in NARC buffer (20 mM HEPES, 400 mM NaCl, 1 mM EDTA) + PI, shaken at 4°C for 30 minutes and centrifuged at 16.000 g for 10 minutes. The supernatant was the nuclear extract.

#### DNase I Hypersensitivity Assay

1x10^6^ cells per condition were transfected with pCMV6 and both *LOC339803* overexpression plasmids. Then cells were washed and collected with cold PBS. Cells were resuspended in C1 lysis buffer (1.28 M sucrose, 40 mM Tris-HCl pH 7.5, 20 mM MgCl_2_, 4 % Triton X-100) and incubated in ice for 15min. Nuclei were pelleted, resuspended in Nuclei wash buffer (10mM Tris pH 7.4, 60mM KCl, 15mM NaCl, 5mM MgCl_2_, 30mM sucrose) and separated in two tubes. One of the tube was incubated with DNase I at 37°C for 30min and the other was left untreated as a negative control. DNA was extracted and same amount of DNA was used to quantify by RT-QPCR. Used primers are listed in Sup Table 2.

#### Bioinformatic packages

MeT-DB V2.0 m^6^A database was used for assessing the existence of m^6^A peaks in *LOC3399803*. MeT-DB V2.0 records predicted transcriptome-wide m^6^A peaks and single-base m^6^A sites from a significantly expanded collection of Methylated RNA Immunoprecipitation Sequencing (MeRIP-Seq) samples. It provides a genome browser to help visualize the m^6^A sites from different studies.

SRAMP (sequence-based N^6^-methyladenosine (m^6^A) modification site predictor) was used to predict m^6^A modification sites on the allele specific RNA sequences of LOC339803 and for secondary structure prediction.

GO MOLECULAR FUNCTION analysis was used with *LOC339803* bound nuclear proteins in HCT-15 intestinal cells from our RIP-MS and p-value results were illustrated using GraphPad Prism 8 (GraphPad Software).

GEPIA (Gene Expression Profiling Interactive Analysis) is an interactive web server for analyzing the RNA sequencing expression data of tumors and normal samples from the TCGA and the GTEx projects. It was used to study the overall survival and expression profiles in GI cancers using *survival analysis* and *boxplots* functions.

## ACKNOWLEDGEMENT

The authors thank for technical and human support provided by general proteomic service from SGIker (UPV/EHU/ ERDF, EU).

This research was supported by Ministerio de Ciencia, Innovación y Universidades grant PGC2018-097573-A-I00 (ACR), Ministerio de Ciencia, Innovación y Universidades grant PID2019-104475GA-I00 (IS), The European Foundation for the Study of Diabetes grant (EFSD)-EFSD/JDRF/Lilly Programme on Type 1 Diabetes Research (IS), Basque Government predoctoral grant PRE_2018_2_0039 (AOG), UPV-EHU predoctoral grant (MSdlC), Ministerio de Ciencia, Innovación y Universidades predoctoral FPI grant PGC2018-097573-A-I00 (HRM).

## DECLARATION OF INTERESTS

The authors declare no competing interests.

## AUTHOR CONTRIBUTIONS

Conceptualization: ACR

Methodology: AOG, HRM, MSdlC, AO, LMMG, IS, ACR

Human sample collection: AHM

Funding acquisition: IS, ACR

Supervision: ACR

Writing – original draft: AOG, ACR

Writing – review & editing: AOG, HRM, MSdlC, AO, LMMG, AHM, IS, ACR

---

**Figure S1.**
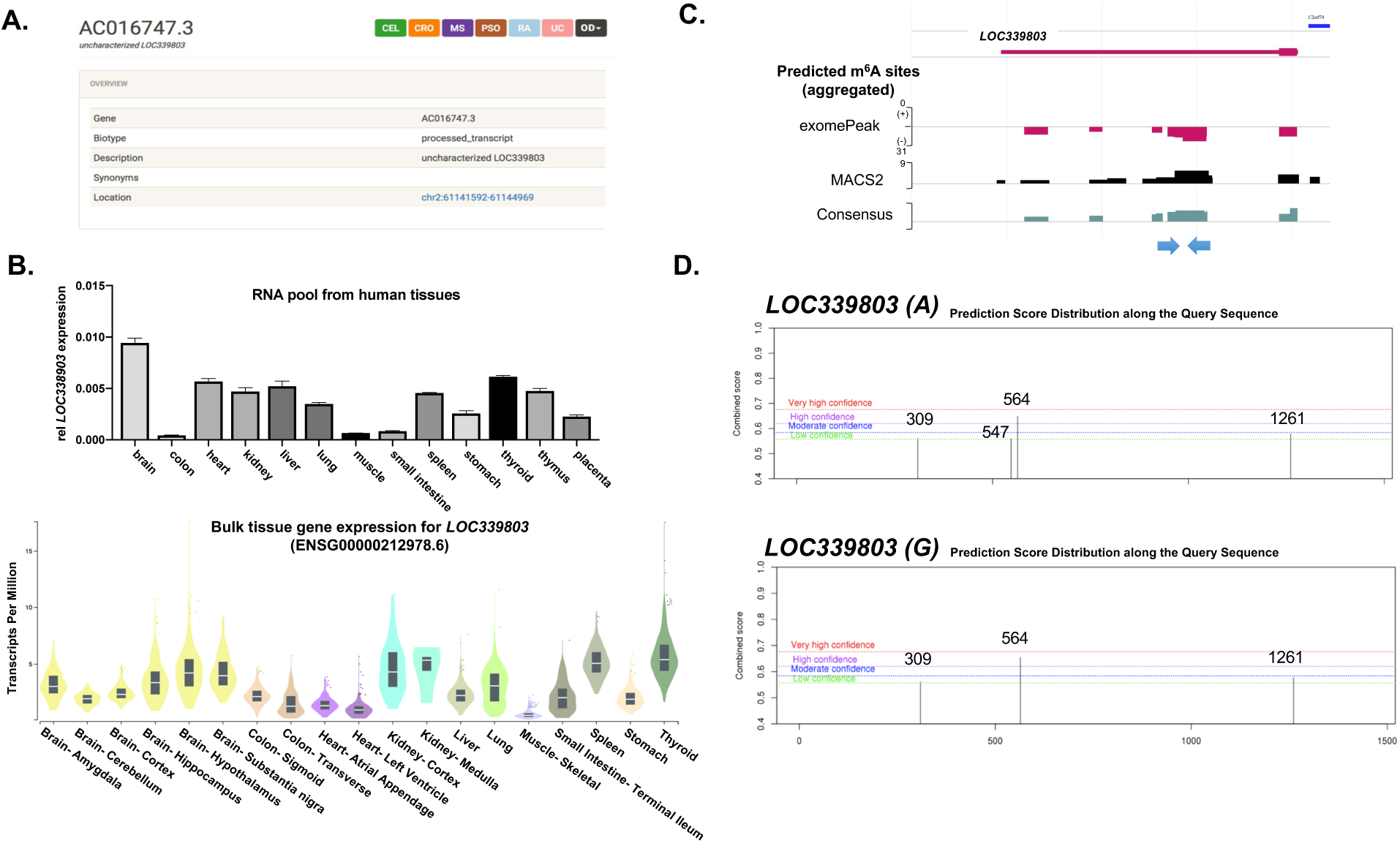
IBD-associated rs11498 SNP genotype affects m^6^A methylation levels in *LOC339803*. **(A)** *LOC339803* (also known as AC016747.3) association to celiac disease (CEL), Chrons’ disease (CRO), multiple sclerosis (MS), psoriasis (PSO), rheumatoid arthritis (RA), ulcerative colitis (UC) according to Immunobase database (https://genetics.opentargets.org/immunobase). **(B)** Up, relative expression of *LOC339803* in commercially available RNA pool of different human tissues, and down, *LOC339803* RNAseq data from GTEx database (https://gtexportal.org/home/). **(C)** m^6^A peaks in *LOC339803* according to MetDB m^6^A database. Arrows indicate the position of the meRIP primers (http://compgenomics.utsa.edu/MeTDB/). **(D)** Allele-specific predicted m^6^A methylation in *LOC339803* according to m^6^A predictor SRAMP online tool (https://www.cuilab.cn/sramp).

**Figure S2.**
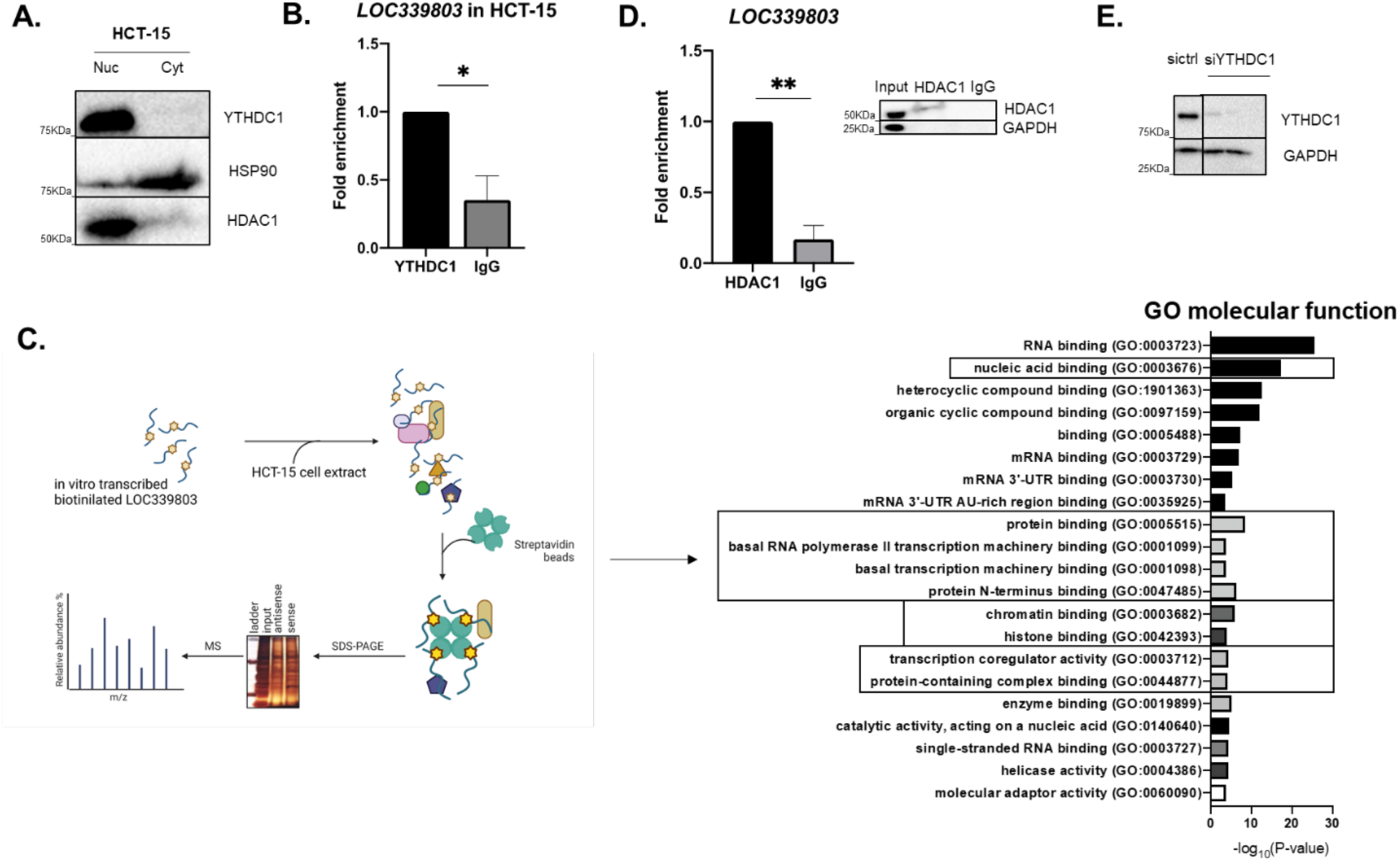
YTHDC1 m^6^A reader influences cellular localization and protein binding of *LOC339803*. **(A)** Protein quantification by western blot in nuclear and cytoplasmic fractions in HCT-15 intestinal cells. HSP90 (cytoplasmic) and HDAC1 (nuclear) were used as controls. **(B)** YTHDC1 immunoprecipitation and quantification of bound *LOC339803* levels assessed by RT-qPCR in HCT-15 intestinal cells. Data are means ± SEM (n=3 independent experiments). **(C)** Left, graphical representation of performed protocol for RIP-MS created with BioRender. Right, GO molecular function enrichment analysis in *LOC339803* binding nuclear proteins from RIP-MS in HCT-15 cells. **(D)** HDAC1 RIP and quantification of bound *LOC339803* levels assessed by RT-qPCR in HCT-15 intestinal cells. Representative immunoblot of the RIP experiment with GAPDH as negative control for the IP. Data are means ± SEM (n=3 independent experiments). **(E)** Representative immunoblot of *YTHDC1* silencing. GAPDH was used as loading control. All p-values determined by one-tailed Student’s t-test. *p<0.05, **p<0.01.

**Figure S3.**
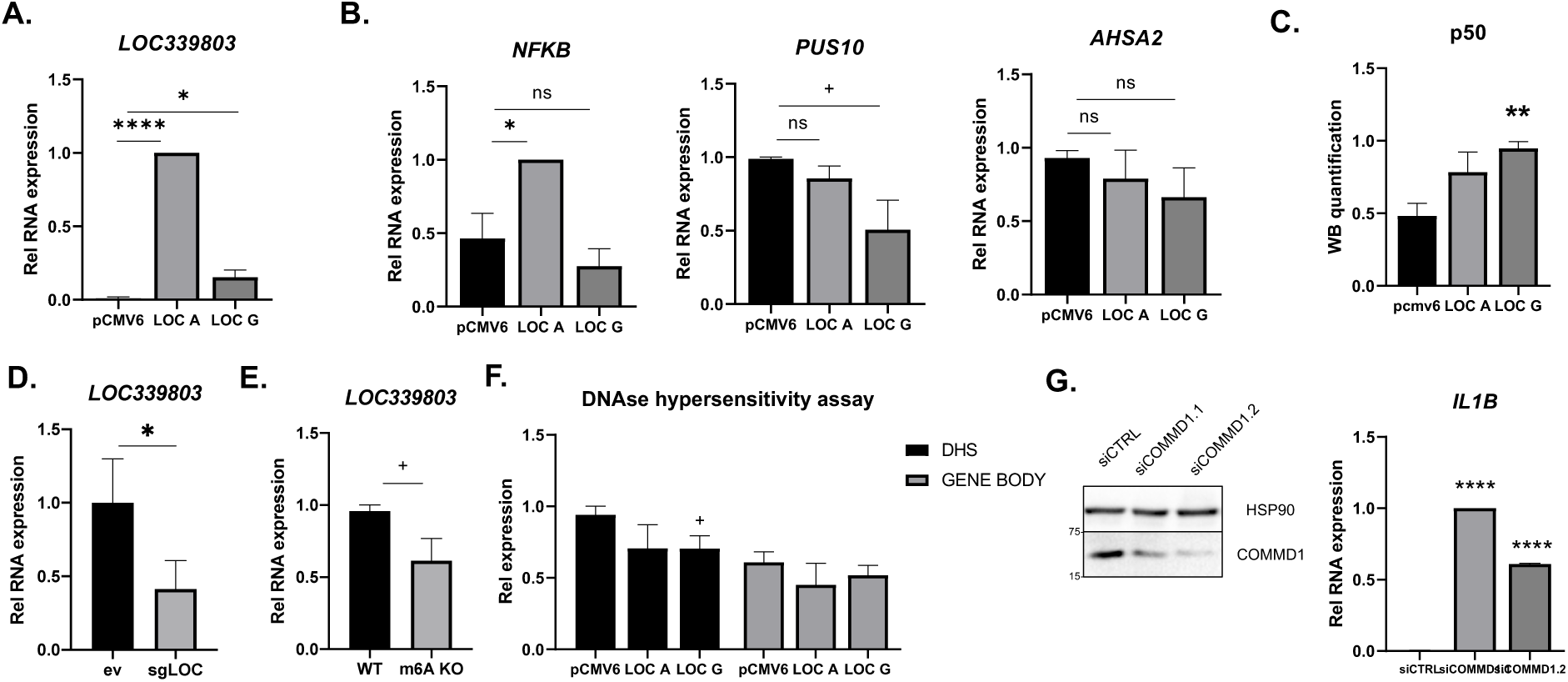
*LOC339803* induction promotes transcriptional repression of *COMMD1* activating NFkB proinflammatory pathway in intestines. Quantification of *LOC339803* (**A**) and (**B**) *NFKB, PUS10* and *AHSA2* RNA levels by RT-qPCR in cells transfected with overexpression plasmids of both forms of *LOC339803* (*LOC A* and *LOC G*). Data are means ± SEM (n=3 independent experiments). p-values determined by two-tailed Student’s t-test. (**C**) Quantification of p50 protein levels in cells transfected with overexpression plasmids of both forms of *LOC339803* (*LOC A* and *LOC G*). Data are means ± SEM (n=4 independent experiments). p-values determined by two-tailed Student’s t-test. (**D**) Quantification of *LOC339803* RNA levels by RT-qPCR in cells with depleted *LOC339803* using CRISPR-Cas9. Data are means ± SEM (n≥5 independent experiments). p-value determined by one-tailed Student’s t-test. (**E**) Quantification of *LOC339803* RNA levels by qPCR in cells with a deletion of the m^6^A methylation region in *LOC339803* using CRISPR-Cas9. Data are means ± SEM (n=4 independent experiments). p-value determined by two-tailed Student’ s t-test. (**F**) Quantification of chromatin accessibility using primers flanking a DNAse I hypersensitibity site (DHS) within *COMMD1* promoter and primers targeting the gene body as control. **(G)** Left, representative immunoblot of *COMMD1* silencing with GAPDH as control. Data are means ± SEM (n=3 independent experiments). p-value determined by two-tailed Student’s t-test. +p<0.1, *p<0.05, **p<0.01, ****p<0.0001.

**Figure S4.**
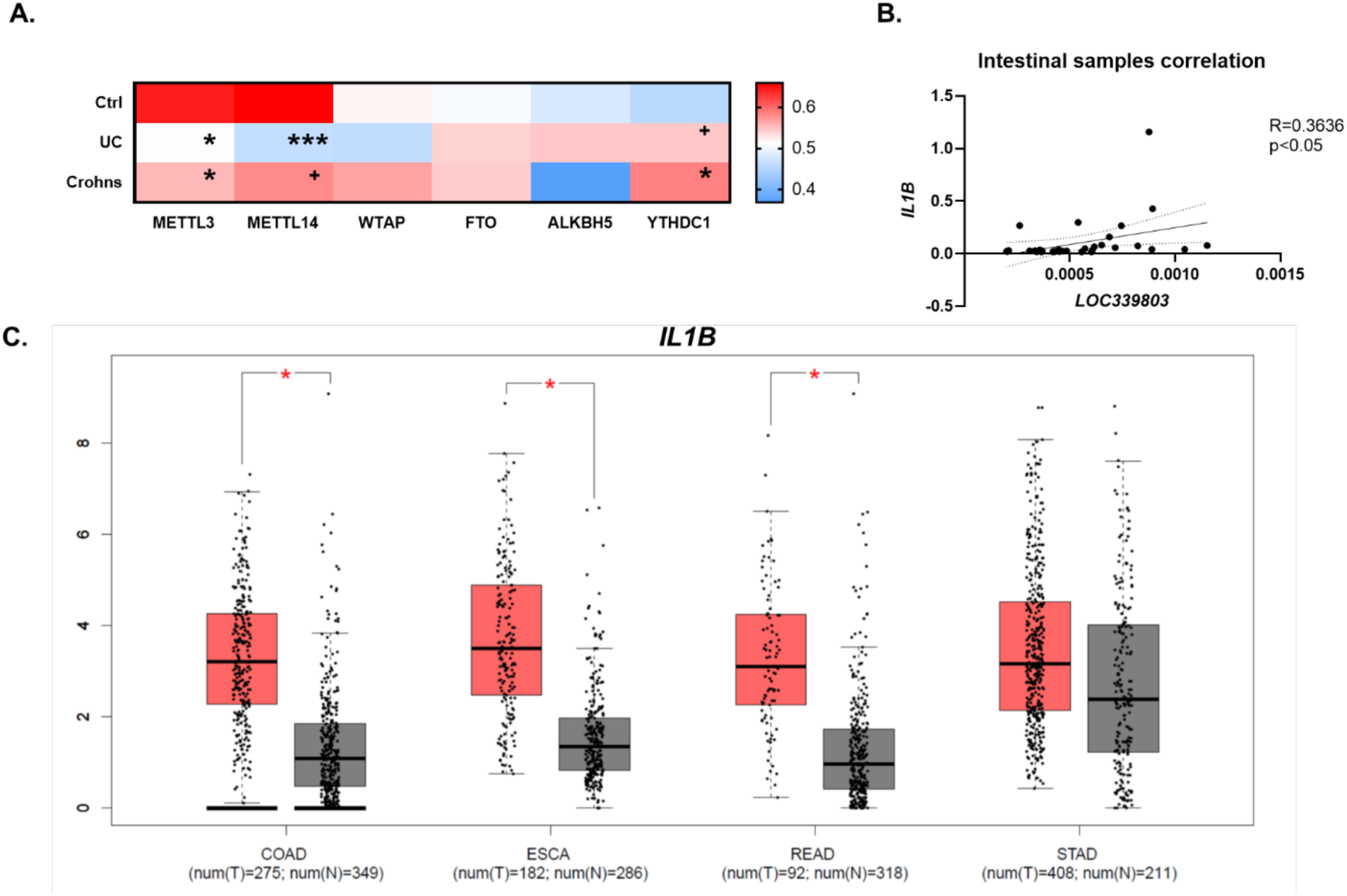
Increased *LOC339803* expression in IBD patients increase risk to develop GI malignancies. **(A)** Heatmap of relative expression of m^6^A machinery genes (*METTL3*, *METTL14*, *WTAP*, *FTO*, *ALKBH5* and *YTHDC1*) quantified by RT-qPCR in intestinal biopsies from controls and patients with ulcerative colitis (UC) and Crohn’s disease (Crohns). Data are represented as mean values (n≥10). **(B)** Correlation between *IL1B* and *LOC339803* in intestinal biopsies. r^2^ and p were calculated by Pearson correlation (n=36). **(C)** *IL1B* expression boxplots for the different GI cancers from GEPIA (http://gepia.cancer-pku.cn/detail.php?gene=&clicktag=boxplot).

**Figure S5.**
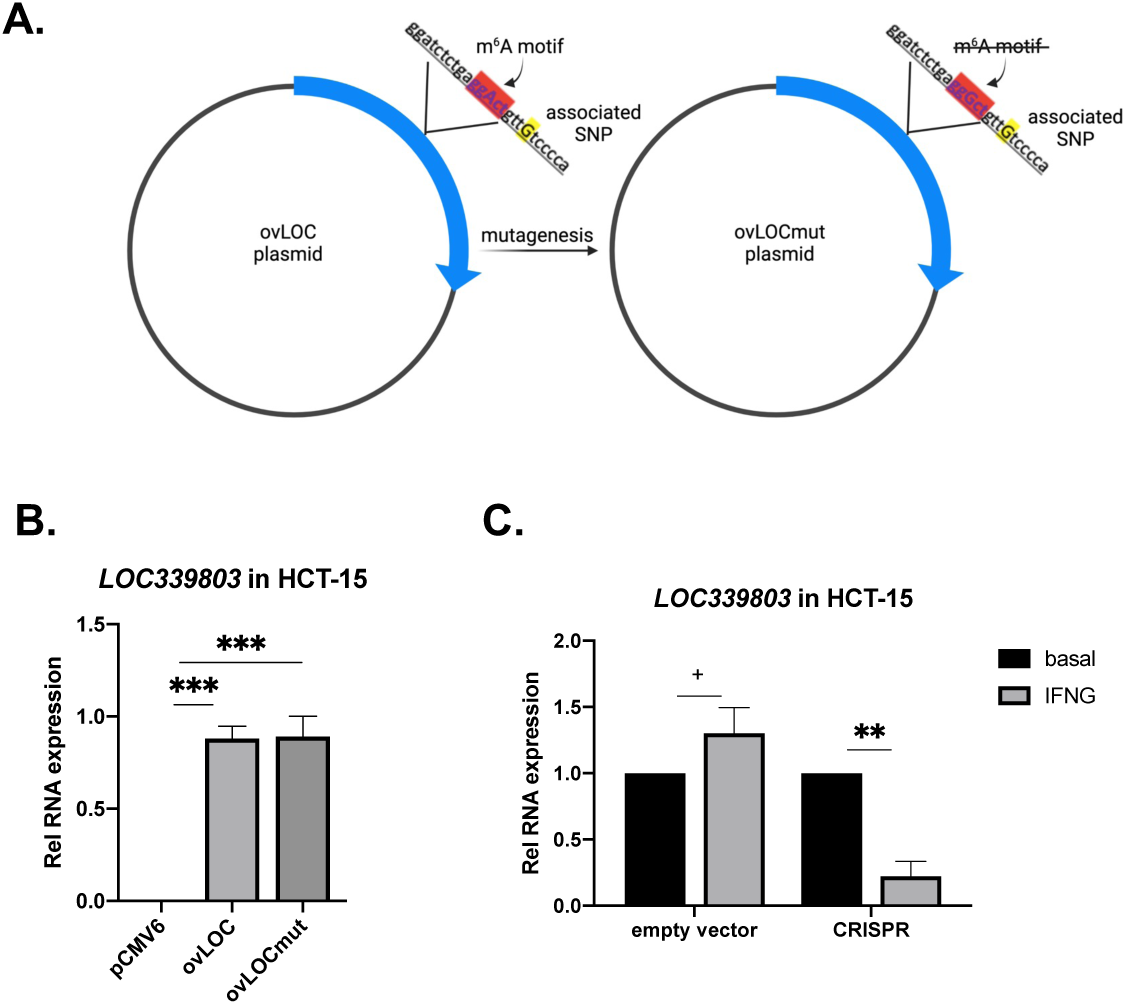
*LOC339803* targeting as a therapeutic strategy. **(A)** Graphical representation of the mutagenesis performed in *LOC339803* overexpressing plasmid to remove m^6^A motif created with BioRender. **(B)** Quantification of *LOC339803* RNA levels by RT-qPCR in cells transfected with an empty vector (pCMV6), *LOC339803* overexpressing plasmid (*ovLOC*) or *LOC339803* overexpressing plasmid with mutated m^6^A motif (*ovLOCmut*) in HCT-15 cells. Data are means ± SEM (n=4 independent experiments). p-values determined by one-way ANOVA test. **(C)** In vitro stimulation model in HCT-15 intestinal cells using interferon gamma (IFNG). Quantification of *LOC339803* RNA levels by qPCR in cells transfected with an empty vector or *LOC339803* depleted cells using CRISPR-Cas9 (sgLOC). Data are means ± SEM (n=3 independent experiments). p-values determined by one-tailed Student’s t-test. +p<0.1, *p<0.05, **p<0.01, ***p<0.001.

**Figure S6.**
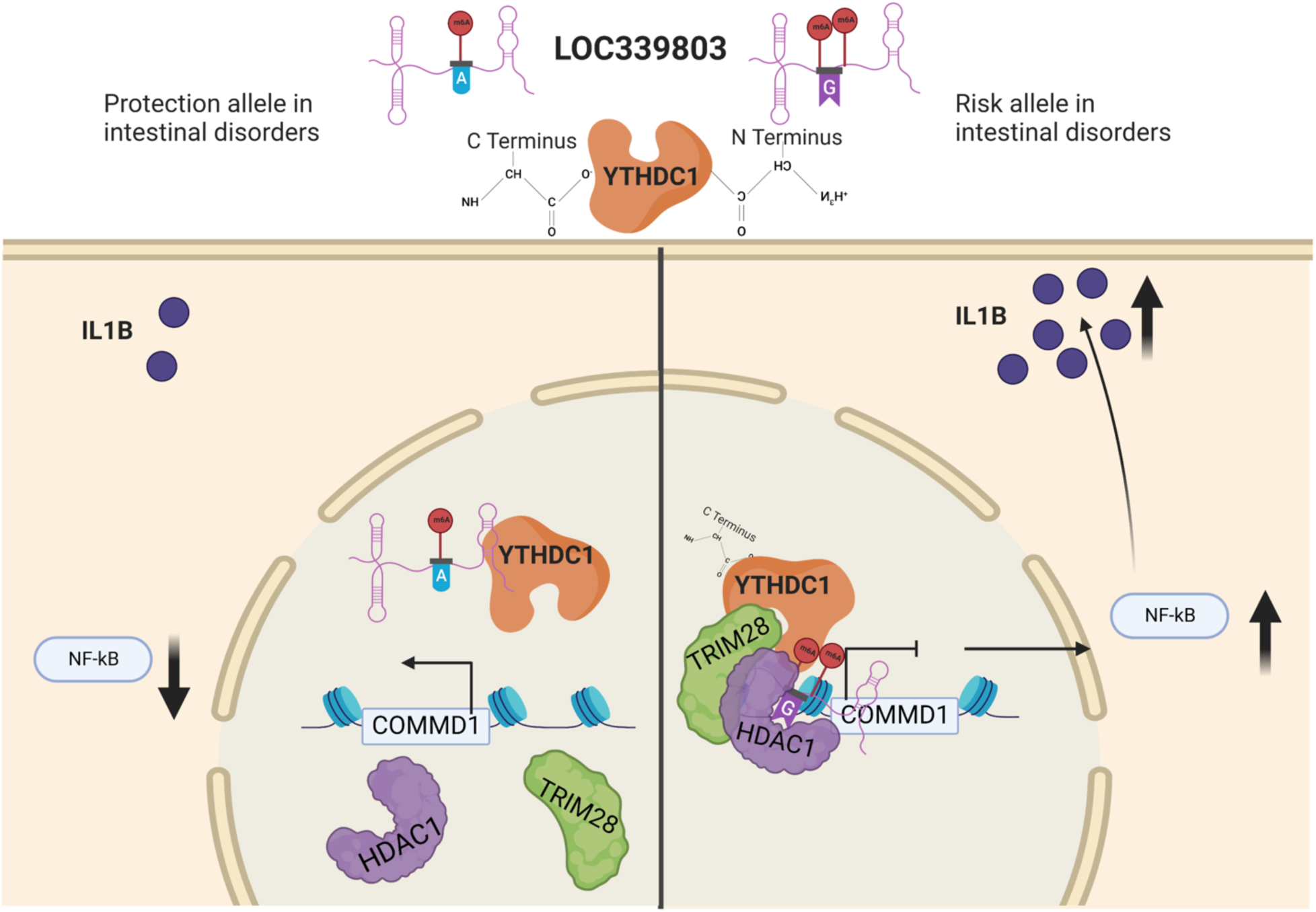
Schematic representation of *LOC339803* mechanism of action.

**Table S1.** List of LOC339803 bound proteins in HCT-15 cell lines derived from RIP-MS. (Separate file)

**Table S2.**
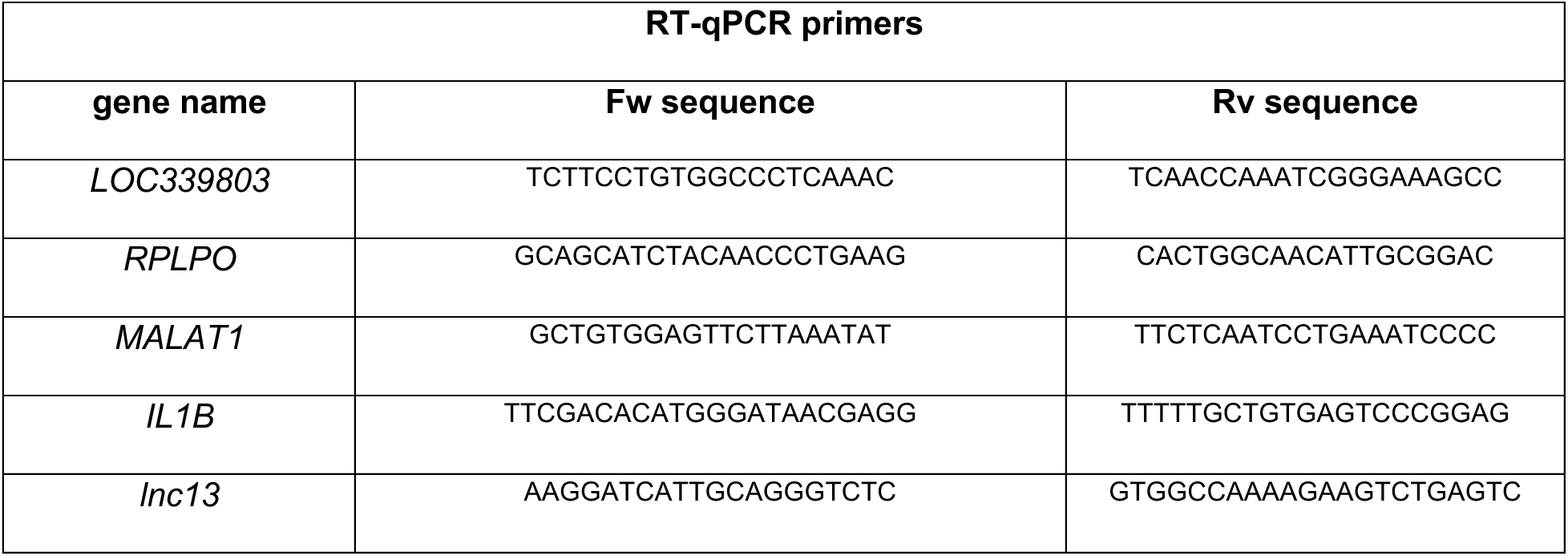
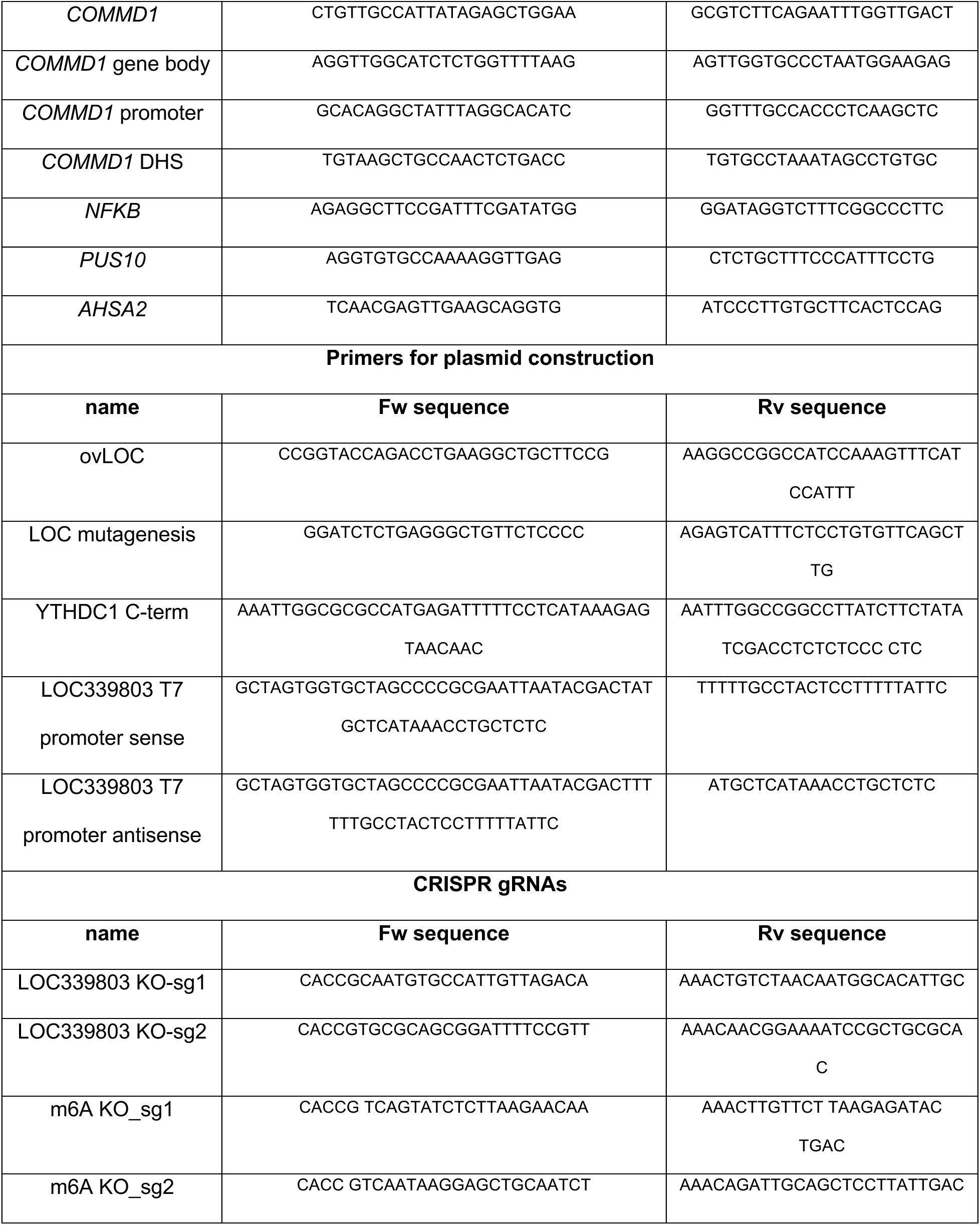
List of primers and their sequences.

## Notes

### Competing Interest Statement

The authors have declared no competing interest.

